# Humans Modulate Walking Speed in Response to the Perceived Energy-Time Costs of Others

**DOI:** 10.64898/2026.07.22.739970

**Authors:** Ryan T. Schroeder, Korbin Allan, Hallee Nugent

**Affiliations:** Yousef Haj-Ahmad Department of Engineering, Brock University, St. Catharines, Ontario, Canada; Department of Kinesiology, Brock University, St. Catharines, Ontario, Canada

**Keywords:** gait biomechanics, locomotion speed, prosocial movement, energy optimization, movement strategy, interpersonal coordination

## Abstract

Humans tend to walk at speeds that minimize energy expenditure and time duration. While most walking experiments examine individuals in relative isolation, everyday locomotion frequently occurs in social contexts. We investigated whether walking speed is modulated in response to a social interaction during a cooperative task. Participants completed 96 randomized walking trials where they approached and retrieved boxes varying in distance (2.5-10 m) and mass (0-6.8 kg). Boxes either rested on the ground or were handed off by an experimenter, signaling prosocial effort benefitting the participant. Approach speed was measured with inertial measurement units placed at the feet and fit to a saturating exponential function of walking distance using a nonlinear mixed-effects regression model. Based on the energy-time optimization framework, we hypothesized that participants would approach more quickly when larger boxes were held at farther distances, to reduce energy and time costs of the experimenter, despite exerting more effort themselves. Participants walked 8.5% faster (1.29 m s^−1^ versus 1.19 m s^−1^; *p* = 3.85 x 10^−6^) when the largest box was held out by the experimenter versus left on the ground. However, the manner in which the box was held had no influence on approach speeds (*p* = 0.31). Exploratory analyses identified modest trends between experiment responses and individual characteristics, but none reached statistical significance. The findings suggest that locomotor decisions reflect not only an individual’s own energy and time costs but also the perceived costs borne by others. This study demonstrates that social context can meaningfully influence walking behavior.

**Summary Statement:** Humans walk more quickly in response to the perceived energy and time others spend helping them, even at the cost of exerting more energy themselves.

## Introduction

Speed is a fundamental characteristic of locomotion that is often recognized for its strong influence on metabolic energy. Longstanding empirical evidence suggests that humans and animals regularly select gait speeds aligned with minimal metabolic energy expenditure per unit distance travelled, commonly referred to as the cost of transport (Bertram, 2005; Bertram & Ruina, 2001; Gutmann et al., 2006; Hoyt & Taylor, 1981; Ralston, 1958; Schmidt-Nielsen, 1972). Such studies have often considered walking and other gaits under steady conditions over relatively long distances or durations. However, real-world walking commonly involves relatively short bouts of movement between discrete locations, with about half of daily walking occurring in under 16 steps at a time (Carlisle & Kuo, 2023; Orendurff, 2008). Recent studies have shown that individuals choose slower speeds when walking over shorter distances, but select faster speeds consistent with the minimum cost of transport at longer distances (Carlisle & Kuo, 2023; Seethapathi & Srinivasan, 2015). The overall gait strategy is well predicted by an energy-time optimization framework (Carlisle & Kuo, 2023), where faster speed reduces the time spent walking but increases energy cost, rising nonlinearly with speed (Bastien et al., 2005). Within this framework, walking speed emerges from a trade-off between energy expenditure and valuation of time.

While the energy-time optimization framework successfully explains locomotor strategies of isolated individuals, it is unknown if it can also explain interactions *between* individuals. In fact, many human locomotion studies examine research participants performing prescribed walking tasks under carefully controlled conditions, and social interactions are intentionally minimized to isolate the effects of interest. Yet, everyday walking often occurs in social contexts, where the presence, actions, and perceived efforts of others may influence locomotor decisions (Huber et al., 2014; Moussaïd et al., 2010; Nicolas & Hassan, 2023).

Even the mere presence of observers have long been known to affect human movement performance, known as social facilitation (Van Meurs et al., 2024). Performance in strength- or endurance-based tasks involving sustained physical effort is generally enhanced by the presence of others; e.g., runners run faster and weightlifters lift more total weight in the presence of observers (Sheridan et al., 2019; Worringham & Messick, 1983). On the other hand, performance in coordination-based tasks requiring precision or fine motor control is sometimes impaired or unaffected by outside observers (Butki, 1994; Krendl et al., 2012; Paulus & Cornelius, 1974; Van Meurs et al., 2024). One explanation for these mixed findings is that social presence competes for limited attentional resources, facilitating relatively automatic tasks while interfering with more attentionally demanding ones (Baron, 1986).

Beyond the effect of a passive observer, social interactions sometimes involve movement coordination which is more interactive. For example, interpersonal coordination refers to dyads or groups of people who spontaneously synchronize walking in real time (Cornejo et al., 2017; Nessler & Gilliland, 2009; Sylos-Labini et al., 2018; Zivotofsky & Hausdorff, 2007). Studies have also shown that upper limb movements are modified to facilitate the actions of others, even while accepting additional effort, during cooperative tasks (Sebanz et al., 2006). Furthermore, prosocial tendencies are observed in individuals who willingly exert extra physical effort in order to benefit others, even when they are strangers (Lockwood et al., 2021). One locomotor example comes from an observational study by Santamaria and Rosenbaum (2011), which identified people approaching a doorway held open, using video data of a public space. The study noted that walking speed may be increased in this context to reduce the total shared effort experienced by both individuals, which they termed a “shared effort” model. In this framework, the approaching individual may willingly expend greater effort individually in order to reduce the effort or waiting time imposed on another person.

While many of these studies demonstrate examples of movement coordination with others, little is known about whether or how locomotor decisions are modified to account for the energy and temporal costs experienced by others during social interactions. Furthermore, while observational studies provide important insight into naturally occurring social behavior, controlled experimental paradigms help support causal inference and facilitate quantitative assessment of resulting biomechanical responses. Therefore, the current objective is to evaluate how humans modify walking speed in response to a controlled social coordination task and contextualize the results within an energy-time optimization framework. To this end, we devised a load-retrieval task in which participants walked to retrieve a box from another person, where the box was either held by the other individual or lay resting on the ground as the individual stood idly by. This paradigm allows walking speed to be examined, not only in the context of the participant’s energy-time costs, but also the perceived energy-time costs of the interacting individual on the participant’s behalf. We hypothesized that participants would approach at faster walking speeds when: (1) the retrieval distance was greater, consistent with previous energy-time optimization results (Carlisle & Kuo, 2023); (2) the box was actively held rather than resting on the ground, thereby signaling energy and time exerted by the holder; (3) the box was larger and more massive, thus increasing the energy cost associated with picking it up and holding it.

## Methods

### Human Participants

A convenience sample of eleven healthy adults (five male, six female) with no known musculoskeletal or neurological impairments affecting gait or the ability to carry a 6.8 kg box was recruited to participate in the study. Participants had a mean ± s.d. height of 1.73 ± 0.12 m, body mass of 77.8 ± 17.8 kg, and age of 26.6 ± 17.8 years. All participants provided written informed consent prior to participation. The study was approved by research ethics boards at the University of Calgary (REB16-1517) and Brock University (REB24-327). All procedures were performed in accordance with the relevant guidelines and regulations.

### Experimental Protocol

All experiments were conducted in a large open space at the Human Performance Laboratory at the University of Calgary. Participants were initially allowed to familiarize themselves with all the boxes in the experiment; they were encouraged to pick them up, feel their weight, and walk around with them, in order to ensure comfortability with the task, and also to become familiar with the effort required to lift them. Following familiarization, participants were asked to stand still at a mark on the ground and await instructions. At the beginning of each trial, the experimenter stood by the appropriate box at a predetermined distance in front of the participant. The experimenter then provided a clear verbal “Go” signal, upon which the participant began walking toward the box, retrieving it from the experimenter (i.e., *Hand off*, Fig. 1A), pausing for a moment, and then returning to the starting mark. The box was then placed on the ground, and the participant awaited the next “Go” signal at the starting mark.

**Figure 1.**
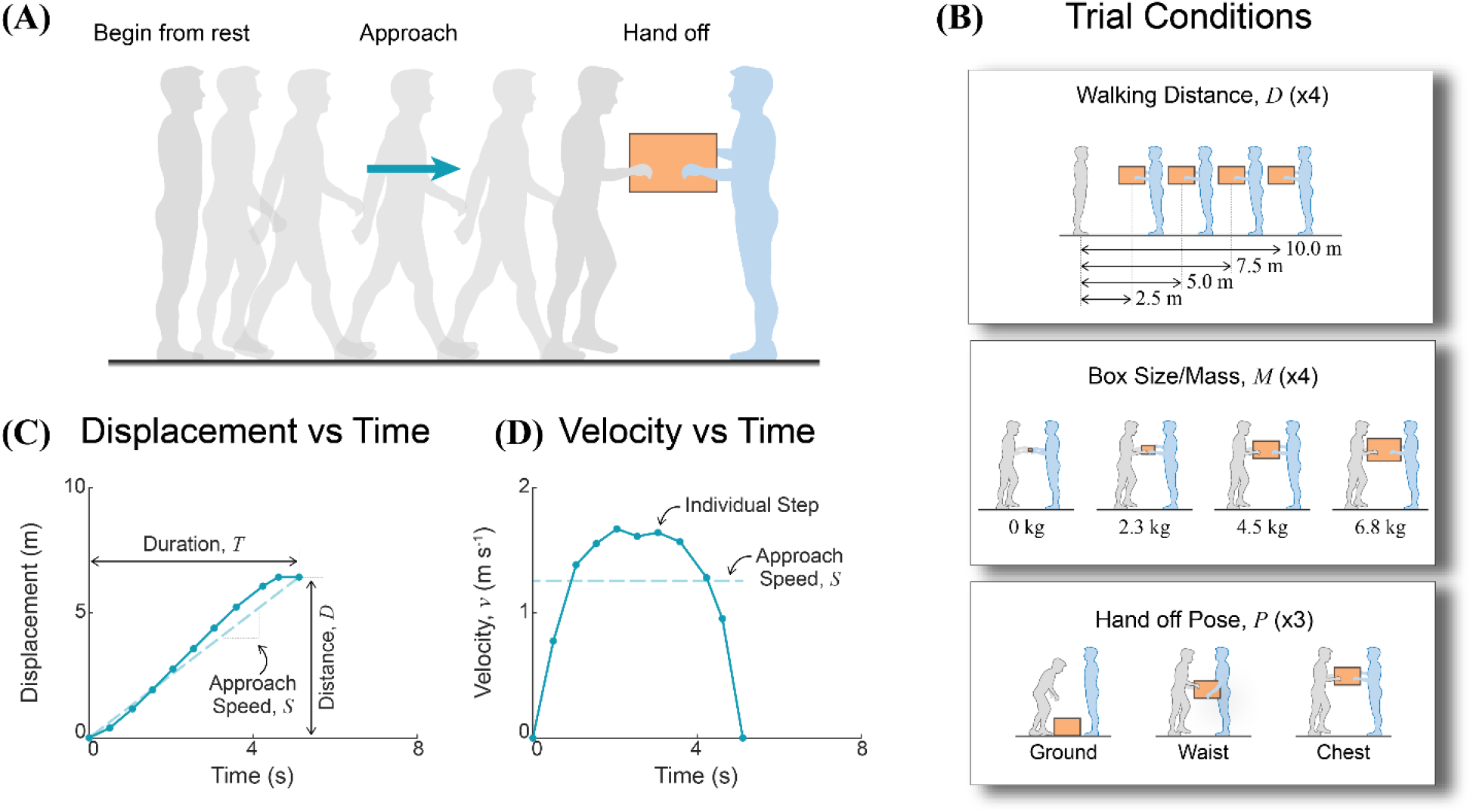
Task, conditions, and example data. **(A)** The task is shown, where the participant begins standing at rest, approaches the box upon receiving a “Go” signal, receives the box during *Hand off*, and then returns the box back to the starting position to await the next trial. **(B)** The experiment trial conditions are shown, where *Walking distance D* is varied across four levels (2.5-10.0 m), *Box size/mass M* is varied across four levels (0-6.8 kg), and *Hand off pose P* is varied across three levels (the box rests on the *Ground*, is picked up and held at *Waist* level, or is picked up and held at *Chest* level with arms outstretched). Example trial data for **(C)** Displacement vs Time and **(D)** Velocity vs Time are shown for an individual trial where discrete data points represent individual steps and time *Duration T*, walking *Distance D*, and *Approach Speed S* are labeled. Trial conditions associated with panes (C) and (D) are: *D* = 7.5 m, *M* = 10 kg, and *P* = Chest.

The above sequence constituted a single trial, which was repeated multiple times in unique combinations of three varied experimental parameters: box distance, box mass, and hand off pose (Fig. 1B). Box distance, defined as the distance from the starting point to the box location in each trial, was varied between 2.5, 5.0, 7.5, and 10.0 m. The box mass was either 0, 2.3, 4.5, or 6.8 kg (0, 5, 10, or 15 lbs weight, respectively). Furthermore, the area of the box face visible to the participant was selected to increase approximately with box mass to help signal increasing load, ranging from 0.088 to 0.248 m². A small hand-sized box was used for the zero-load condition, whereas the other boxes contained compacted sandbags secured at the bottom to reduce shifting while being carried. The weight of each box was explicitly labeled so that participants knew the loads even before the first walking trial. The pose of the experimenter during hand off was also varied. Either the box rested on the ground as the experimenter stood still nearby, or it was picked up from the ground and held at waist level, or it was picked up from the ground and held at chest level with arms outstretched toward the participant. These poses were chosen to represent increasing levels of effort by the experimenter on behalf of the participant, in order from least to most effort: ground, waist, and chest level.

Each unique combination of experimental parameters (4 x 4 x 3 = 48 conditions) was repeated twice for a total of 96 walking trials per data collection with a participant. Trial conditions were randomized to reduce ordering effects. Each trial was timed to last one minute, regardless of how soon the participant returned to their starting point, and participants were informed of the time schedule at the outset to eliminate participant incentives to rush through the protocol and finish early. Participants were instructed to walk and carry the loads as naturally as possible. They were told that the purpose of the study was to investigate how people carry loads and were not informed that social interaction was a primary manipulation in the experiment. During all trials and sessions, the same male experimenter performed the box hand off with all participants. They aimed to maintain a consistent, neutral demeanor, avoid displaying overt visible signs of struggle while picking up or holding the boxes, and minimize unnecessary social interaction with participants as much as possible.

Immediately following completion of the walking trials, participants completed an online questionnaire, administered using Qualtrics (Qualtrics, Provo, UT, USA), based on the Toront Empathy Questionnaire (Spreng et al., 2009). The questionnaire was used to examine potential relationships between empathy scores and participant responses during the experiment.

### Instrumentation and Analysis

Inertial Measurement Units or IMUs (Opal V2R, APDM Wearable Technologies, Portland, OR, USA) were secured to the participants’ shoes near the toe and recorded data at an acquisition rate of 128 Hz. Additional IMUs were strapped to the wrists of the participants, the wrist of the experimenter, and the top of the box used in each trial. The wrist and box IMUs were not formally analyzed but helped verify trial timing, box hand off events, and condition identification during analysis. Additionally, the experimenter used their wrist-mounted IMU to create timestamps corresponding to the verbal “Go” signal, allowing the beginning of each trial to be identified quantitatively. The end of the approach phase was determined by the final footfall prior to box retrieval and thus did not include the time required to pick up or retrieve the box after the last step had been taken.

IMU data were analyzed following the protocol described by Rebula et al. (2013). Briefly, data were segmented into individual strides using periods of near-zero rotational velocity and acceleration corresponding to foot contacts with the ground. Velocity and vertical position of the foot was reset to zero at each of these points, assuming flat ground, and to help mitigate excessive drift error. Accelerations were then integrated twice to determine foot displacements within each stride, assuming zero initial acceleration and velocity conditions at the beginning of each stride in stance. Strides were converted into steps by compiling kinematic data from both feet. Specifically, fore-aft displacement of the foot was calculated from the time of its contact to the subsequent contralateral foot contact giving step length, while step time was the time interval between contacts. Step velocity was then calculated as step length over step time and was considered an approximate indication of center-of-mass velocity over time.

Overall displacement of the body was defined as the net displacement of a series of step lengths and plotted as a trajectory in time (Fig. 1C). Step velocity was additionally plotted as a profile in time (Fig. 1D). It was assumed that individuals walked approximately in a straight line, and thus, the trajectories characterized progress in the overall direction of travel. Velocity profiles associated with individual trials were time normalized, interpolated at common intervals, and averaged at the corresponding values. The results were then rescaled by the mean time duration in order to produce mean profiles across all participants and/or trials associated with a given set of experimental parameters (e.g., Fig. 2A, thick lines in inset plot). During interpolation, the number of intervals was chosen to match the trial with the fewest data points included in the mean, to avoid oversampling.

**Figure 2.**
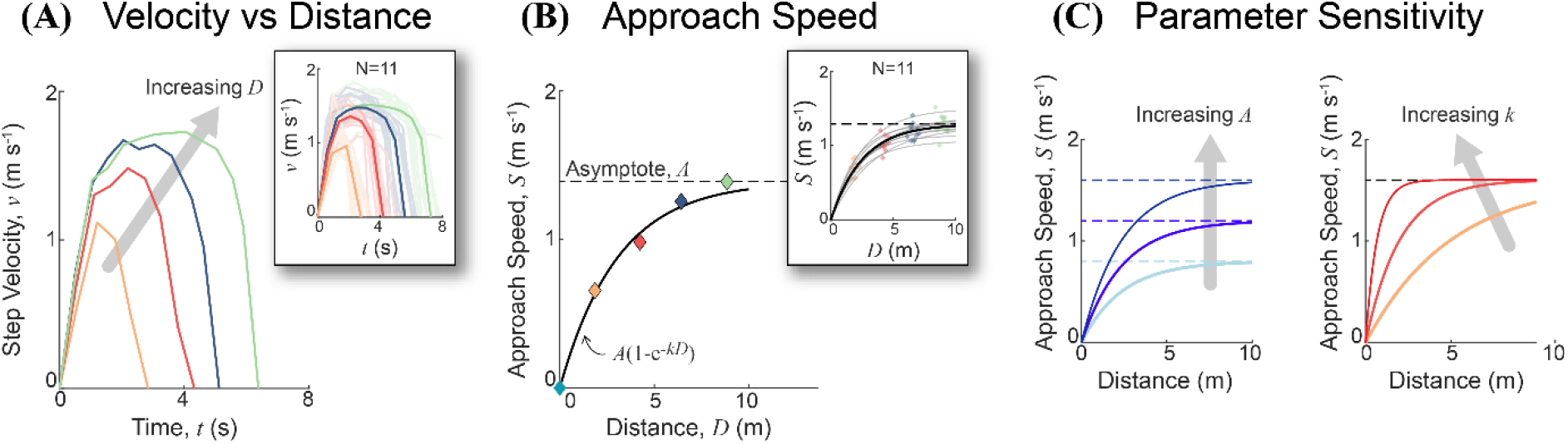
Speed changes with walking distance. **(A)** Step velocity profiles over time are shown for a representative participant for all tested distances (distinguished by color) when approaching a 4.5 kg box held at waist level. In addition, profiles for all participants and the same conditions are shown in the inset plot (N=11, thin lines) as well as the mean for each distance condition (thick lines). **(B)** Overall *Approach Speed S* is shown versus walking *Distance D*, with color of individual data points indicating the corresponding *step velocity* profiles in (A). The black curve indicates the best fit curve of a saturating exponential function, *S* = *A*(1 - *e*^−*kD*^), used to characterize the *Approach-Speed-Distance* relationship. The resulting approach speed *asymptote A* from the regression is plotted as a thin, horizontal dashed line. Similar to (A), the inset contains the data and regression curves for all participants (N=11). **(C)** Sensitivity of the saturating exponential function to the two fitting parameters are shown, where *asymptote A* and *rate constant k* are varied over a range of values (left: *A* = 0.8, 1.2, and 1.6 m s^−1^ and *k* = 0.425 m^−1^; right: *A* = 1.6 m s^−1^ and *k* = 0.2, 0.5, and 1.5 m^−1^).

The approach speed *S* was calculated by dividing the net fore-aft displacement of the feet during approach (walking distance *D*) by the corresponding time duration *T* (Fig. 1C,D). Duration was determined as the time interval between the experimenter’s “Go” timestamp and the completion time of the final footfall prior to box retrieval during hand off. The approach included all steps from the beginning mark to the second to last step before reaching the box. The last step was excluded from analysis since it marked a task transition from approach to hand off, where details of the hand off pose could interfere with the main effect of interest (e.g., individuals may take more time to pick up a box from the ground rather than receive it standing up from the experimenter, regardless of their overall urgency to reach the general box location). Peak approach speed was also recorded in order to draw comparisons with previous studies (Carlisle & Kuo, 2023). However, average approach speed was primarily used in analysis, instead of peak speed, to account for the whole walking bout during approach and to reduce potential variance associated with choosing a single data point per trial. Data associated with participants returning to the starting location carrying the box were not analyzed.

### Statistical Models

Mixed-effects nonlinear regression models were used to assess the effects of the experimental parameters on approach speed while accounting for repeated measurements within participants. Models were fit using the nlmefit() function in MATLAB R2026a (MathWorks Inc., Natick, MA, USA) with default options such as a constant residual error model, LME likelihood approximation, and an unconstrained random-effects covariance matrix. Following Carlisle and Kuo (2023), approach speed *S* was modeled as a saturating exponential function of walking distance *D* (Fig. 2B),

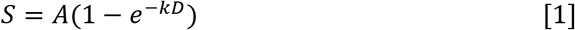

where *A* represents the asymptotic approach speed at long walking distances and *k* is the rate constant describing how steeply the curve nears the asymptote. As such, a higher or lower *A* value is associated with a higher or lower speed asymptote at long distances and a higher or lower *k* value is associated with a steeper or more gradual convergence with the asymptote, respectively (Fig. 2C). Both *A* and *k* were modeled as linear functions of the experimental parameters:

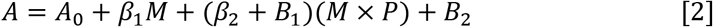

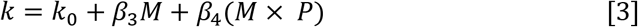

where *A_0_* and *k_0_* are fixed-effect offsets representing nominal walking speeds of the sample when approaching a zero-load box resting on the ground. Additionally, *M* is box mass (in kg) and *P* is hand off pose, a categorical variable describing whether the box was held during hand-off, with the ground condition serving as the reference. The coefficients *β_i_* are fixed effect parameters and *B_i_* are participant-specific random effects. Specifically, parameters *β_1_* and *β_3_* quantify the effects of box mass on approach speed. Parameters *β_2_* and *β_4_*quantify the interaction effect of box mass times pose, indicating whether it was held or left resting on the ground. This interaction term captures the essential influence of the social interaction on walking behavior in the experiment – that is, the degree to which box mass affected participant gait, depending on whether it was held or not. *B*_1_ is used to modify *β*_2_ in order to characterize individual responses to a box being held, while *B*_2_ allows for participants to have individual speed asymptotes in the model.

A variation of the model was also produced, where three levels of the pose variable were used to distinguish effects between boxes held at waist level versus chest level with arms outstretched, relative to the box resting on the ground. A *post hoc* two-sided t-test was conducted to evaluate a statistical difference between the regression coefficients on holding the box at waist level versus at chest level using the ratio of the difference to its standard error.

Model fit was assessed using Akaike’s Information Criterion or AIC (Akaike, 1974), and goodness of fit using root mean squared error or RMSE, as well as coefficient of determination. Fixed-effect parameter estimates were reported alongside 95% confidence limits (CL) and were tested against the null hypothesis that the corresponding regression coefficient was equal to zero. Statistical significance was assessed using an alpha level of 0.05, with Bonferroni corrections (Dunn, 1961) applied to account for multiple hypothesis testing based on the number of parameters evaluated in the model. In addition to the mixed-effects models, separate nonlinear regressions were performed for all participants and experimental conditions to generate individual speed-distance relationships.

Further exploratory, *post hoc* analysis was conducted in order to evaluate any potential relationships between participant-specific responses in the experiment (*β*_2_ + *B*_1_) and individual characteristics such as empathy questionnaire scores, sex, age, and nominal walking speed (i.e., approach speeds associated with a zero-load box resting on the ground). Pearson correlation coefficients and corresponding t-tests were used to quantify any potential relationships between these data. Boxplots were used to visualize data distributions over various experimental conditions, where whiskers are the data range, boxes are interquartile range, and the middle ticks are median values. Outliers were identified using the interquartile range rule (Tukey, 1977) and indicated with + symbols, but were still included in all data analysis and the statistical models.

### Empathy Questionnaire

Following completion of the walking trials, participants completed an online survey based on the Toronto Empathy Questionnaire (Spreng et al., 2009), a self-report measure consisting of 16 statements designed to assess empathic tendencies (e.g., “It upsets me to see someone being treated disrespectfully”). Responses were scored on a five-point scale, with *Never* assigned a score of 0, *Rarely* assigned a score of 1, *Sometimes* assigned a score of 2, *Often* assigned a score of 3, and *Always* assigned a score of 4. Questions alternated between empathy-affirming and empathy-disaffirming statements, with scores reversed as appropriate when calculating the final composite score. Individual responses were summed to obtain a composite empathy score ranging from 0 (least empathetic) to 64 (most empathetic). All questionnaires were completed immediately following the experimental protocol, and no survey responses were missing.

## Results

### Approach speeds asymptote at long walking distances

In a typical trial, participants approached the box with step velocity profiles that followed a characteristic inverted U-shaped pattern. Acceleration occurred over the first few steps, followed by near-steady walking, and subsequent deceleration in the last few steps before hand off (Fig. 2A). For trials with farther walking distances, participants typically employed faster walking speeds, even as duration of the walking bout also increased. Overall, the approach speed increased nonlinearly with walking distance, plateauing toward an asymptote at long distances (Fig. 2B). A saturating exponential function (Eqn. 1) was used to characterize the relationship between approach speed and walking distance (Fig. 2B). The asymptote (*A*) of the function indicates the walking speed converged upon at long distances, while the rate constant (*k*) characterizes how steeply the curve converges on the asymptote. Larger *k* values indicate steeper curves before plateauing, while smaller values indicate a more gradual convergence with the asymptote (Fig. 2C). Nominal approach speed (i.e., the average speeds participants used to approach a small, zero-load box resting on the ground) was characterized by speed asymptote *A_0_* = 1.25 m s^−1^ (CL: 1.18, 1.32 m s^−1^) and rate constant *k_0_* = 0.44 m^−1^ (0.42, 0.45 m^−1^). Greater than 99% of variance was explained by the saturating exponential function in the nominal condition (Eqn. 1, *R*^2^ = 0.993). In fact, nonlinear regression curves conducted on speed-distance data for all participants in all experimental conditions (N=132) resulted in adjusted coefficients of determination ranging from *R*^2^ = 0.961 - 0.999, with an average of *R*^2^ = 0.994. Furthermore, the nonlinear mixed-effects regression model had an AIC value of −1485.8 and a root mean square error of 0.054 m s^−1^, or about 4.3% error of the nominal speed asymptote. Together, these results seem to indicate that the saturating exponential function characterizes speed-distance data reasonably well in the experiment.

### Walking duration is *shorter* when a large box is held at distance

Although time duration and approach speed increased with walking distance, they were also influenced by whether the experimenter held the box or left it to rest on the ground, and the effect magnitude depended on box mass. For example, the experimenter holding a zero-load box produced little to no change in walking duration (−0.31 s or −4.12%) or approach speed (+0.01 m s^−1^ or +0.64%) when compared to trials where the box was left on the ground, for a 10 m walking distance (Fig. 3, left plots, duration changes represented with blue arrows). In contrast, trials with the large box (6.8 kg) induced small to moderate decreases in duration and increases in speed (duration: −0.67 s or −8.36%; speed: +0.05 m s^−1^ or +4.08%). Furthermore, walking distance was reduced by about 0.36 m on average, regardless of box mass and distance, likely due to trials where the box was held out towards participants with arms outstretched (Fig. 3, blue arrows pointing left). Despite the slightly reduced walking distances, duration was sometimes reduced even more (Fig. 3, blue arrows pointing down), ultimately resulting in modest approach speed increases with larger boxes (Fig. 3, right plots). This effect was largely contained to farther walking distances, and in fact, at shorter distances, approach speed was *reduced*, even when approaching a large box.

**Figure 3.**
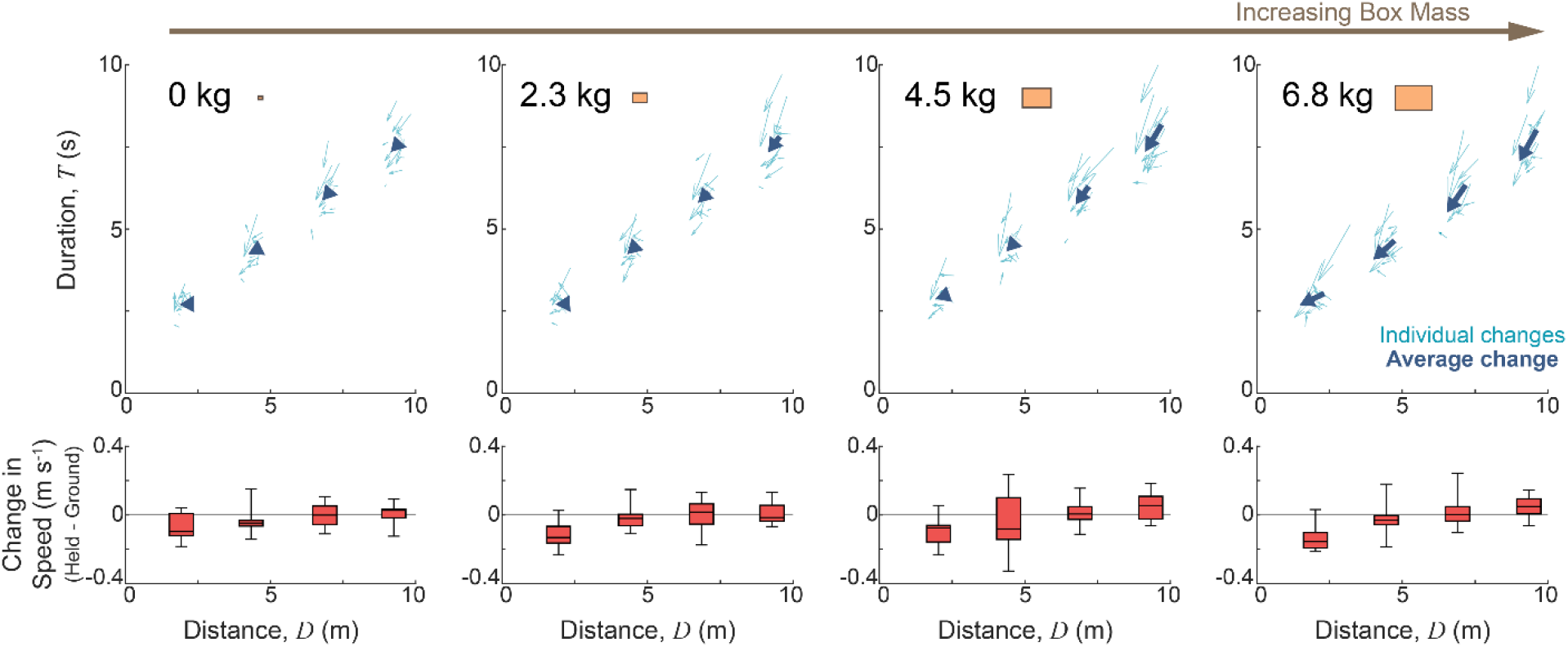
Gait changes in response to the box being held. In the upper panels, time *Duration T* is plotted versus walking *Distance D* using the quiver() function in MATLAB to represent changes in these parameters with hand off pose. Specifically, arrow tails indicate the parameters corresponding to trials where the box rested on the ground and arrow heads indicate parameters corresponding to trials where the box was picked up and held before hand off. As such, the arrow represents a change in *Duration* and *Distance* between trial conditions. Light blue arrows represent individual changes (N=176) while dark blue arrows represent mean changes in each condition. Each grouping of arrows represents a given walking *Distance*, Existing optimization theories explain locomotor decisions using costs borne by the individual walker. Whether humans also incorporate costs experienced by interacting partners remains largely unknown plots left to right represent trials with increasing box size. The lower panels of boxplots show changes in approach speed, where positive values indicate a faster walking speed when the box was held versus when it sat on the ground. The boxplots themselves summarize the range, interquartile range, and median values of all subjects in each condition.

Despite the changes illustrated in Figure 3, direct comparisons of approach speed are difficult to interpret due to variations in walking distance. As shown in Figure 2, average walking speed depends strongly on walking distance, and thus, it is important to account for distance when evaluating speed. To better isolate systematic changes in walking behavior across conditions, comparisons were made using the parameters of the speed-distance relationship (Eqn. 1), which directly account for the effect of walking distance.

### Approach speed is *faster* when a large box is held at distance

For the zero-load box, approach speed was largely unchanged whether the box was held by the experimenter or left on the ground, as evidenced by the near-complete overlap of the corresponding speed-distance data (Fig. 4, left plot). As box mass increased, however, the data began to diverge, particularly at farther walking distances where walking speeds approached the asymptote (Fig. 4, right plot). For the 6.8 kg box, this separation was most pronounced, with higher approach speeds observed when the box was held rather than left on the ground. Overall, systematic trends were observed, where the asymptote *A* was increased from 1.23 to 1.27 m s^−1^ (2.75% increase) for the 2.3 kg box, from 1.21-1.28 m s^−1^ (5.58% increase) for the 4.5 kg box, and from 1.19 to 1.29 m s^−1^ (8.50% increase) for the 6.8 kg box, when the box was held versus left on the ground.

**Figure 4.**
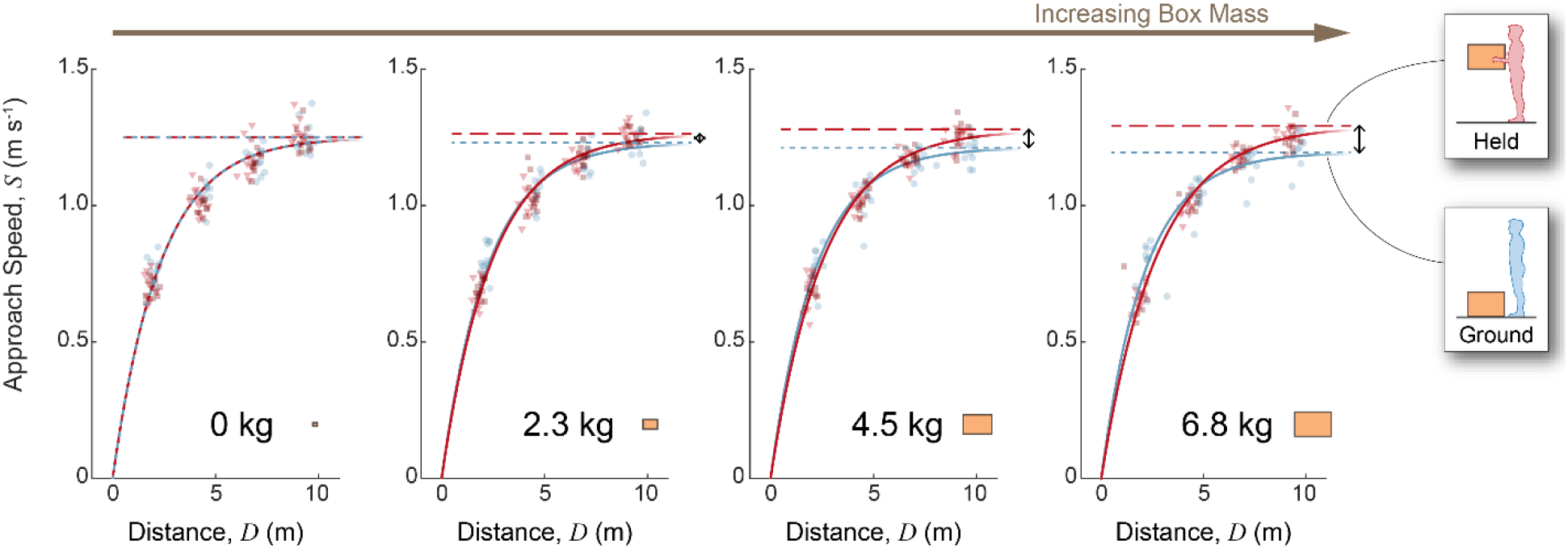
Approach Speed versus walking Distance. Each panel indicates changes in Approach Speed *S* with walking Distance *D*, for each box mass (increasing in mass from left to right). Discrete data points represent individual trials (N=528) for all participants. Light blue circles are trials where the box lay on the ground, while red squares and triangles are trials where the box was held at waist or chest level, respectively. Best fit curves are shown from the nonlinear mixed effects regression model with dashed lines indicating speed asymptotes at long walking distances. The black double-sided arrows show separation between asymptotes of the held and ground curves, increasing with box mass.

Model-derived parameter estimates revealed that the speed asymptote *A* actually *decreased* slightly with box mass when the box was left on the ground (*β*_1_ = −8.46 x 10^−3^ m s^−1^ kg^−1^, *p* = 1.27 x 10^−4*^) but *increased* with box mass when it was held versus left on the ground (*β*_2_ = 1.49 x 10^−2^ m s^−1^ kg^−1^, *p* = 3.85 x 10^−6*^; see Fig. 5A and Table 1). In contrast, the rate constant *k* held the opposite pattern, increasing slightly, albeit not significantly, when the box was left on the ground (*β*_3_ = 5.81 x 10^−3^ m^−1^, *p* = 0.057) and decreasing with mass when the box was held (*β*_4_ = −1.53 x 10^−2^ m^−1^, *p* = 1.73 x 10^−7*^; see Fig. 5B and Table 1). Together, these findings indicate that larger held boxes *increased* approach speeds at *farther* walking distances, but *reduced* approach speeds at *shorter* distances.

**Figure 5.**
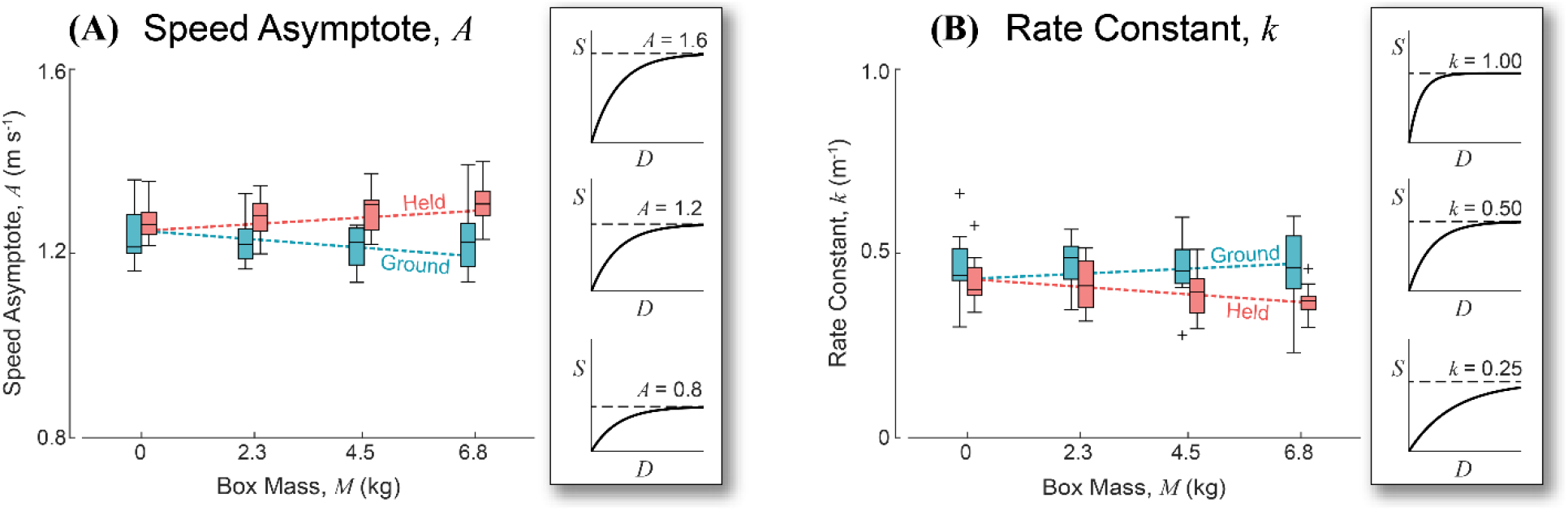
Regression parameter trends across conditions. **(A)** Approach *Speed Asymptote A* is shown to increase or decrease with *Box Mass M*, depending on whether the box was held (red; *β*_2_ = 1.49 x 10^−2^ m s^−1^ kg^−1^, *p* = 3.85 x 10^−6*^) or resting on the ground (blue; *β*_1_ = −8.46 x 10^−3^ m s^−1^ kg^−1^, *p* = 1.27 x 10^−4*^). Inset plots illustrate speed-distance curves with varying *Speed Asymptote* values, for reference. **(B)** *Rate Constant k* shows the opposite trend from *Speed Asymptote*, where values increase when the box was on the ground (blue; *β*_3_ = 5.81 x 10^−3^ m^−1^, *p* = 0.057) and decrease when the box was held (red; *β*_4_ = −1.53 x 10^−2^ m^−1^, *p* = 1.73 x 10^−7*^). Inset plots illustrate speed-distance curves with varying *Rate Constant* values, for reference. In both panes, boxplots display the range, interquartile range, and median of the data (N=88) for all subjects in each condition (+ symbols indicate outlier data). Dashed lines show the effect of *Box Mass M* on each parameter from the nonlinear mixed effects regression model.

**Table 1:** Summary of Fixed Effects in Nonlinear Regression Model Nonlinear mixed effects model fit to 528 observations from 11 individuals (AIC = −1485.8; RMSE = 0.054 m s^−1^). Fixed effects on speed asymptote include nominal asymptote *A*_0_ (speed asymptote when approaching a zero-load box on the ground), box mass *β*_1_ (m s^−1^ kg^−1^), and an interaction between box mass and hand off pose *β*_2_ (m s^−1^ kg^−1^). Fixed effects on rate constant include nominal rate *k*_0_, box mass *β*_3_ (m^−1^ kg^−1^), and an interaction between box mass and hand off pose *β*_4_ (m^−1^ kg^−1^). The fixed effects contribute to speed asymptote *A* and rate constant *k* in the saturating exponential function to describe how approach speed *S* changes with walking distance *D* (see Eqn. 1). Best fit coefficient values (*β*), 95% confidence limits (CL), and *p*-values are shown, marked with asterisks to indicate significance after controlling for multiple testing.

| Description | Fixed Effects |  |  |  |  |
| --- | --- | --- | --- | --- | --- |
|  | Parameters | Estimate | Lower CL | Upper CL | *p* |
| Nominal asymptote | $A_0$ (m s<sup>-1</sup>) | 1.251 | 1.181 | 1.321 | <1.000 x10<sup>-6*</sup> |
| Box mass on asymptote | $\beta_1$ (m s<sup>-1</sup> kg<sup>-1</sup>) | -0.008 | -0.013 | -0.004 | 1.27 x10<sup>-4*</sup> |
| Box-pose on asymptote | $\beta_2$ (m s<sup>-1</sup> kg<sup>-1</sup>) | 0.015 | 0.009 | 0.021 | 3.85 x10<sup>-6*</sup> |
| Nominal rate constant | $k_0$ (m<sup>-1</sup>) | 0.435 | 0.417 | 0.453 | <1.00 x10<sup>-6*</sup> |
| Box mass on rate constant | $\beta_3$ (m<sup>-1</sup> kg<sup>-1</sup>) | 0.006 | 0.000 | 0.012 | 5.72 x10<sup>-2</sup> |
| Box-pose on rate constant | $\beta_4$ (m<sup>-1</sup> kg<sup>-1</sup>) | -0.015 | -0.021 | -0.010 | 1.73 x10<sup>-7*</sup> |

Nonlinear mixed effects model fit to 528 observations from 11 individuals (AIC = -1485.8; RMSE = 0.054 m s<sup>-1</sup>). Fixed effects on speed asymptote include nominal asymptote $A_0$ (speed asymptote when approaching a zero-load box on the ground), box mass $\beta_1$ (m s<sup>-1</sup> kg<sup>-1</sup>), and an interaction between box mass and hand off pose $\beta_2$ (m s<sup>-1</sup> kg<sup>-1</sup>). Fixed effects on rate constant include nominal rate $k_0$ , box mass $\beta_3$ (m<sup>-1</sup> kg<sup>-1</sup>), and an interaction between box mass and hand off pose $\beta_4$ (m<sup>-1</sup> kg<sup>-1</sup>). The fixed effects contribute to speed asymptote $A$ and rate constant $k$ in the saturating exponential function to describe how approach speed $S$ changes with walking distance $D$ (see Eqn. 1). Best fit coefficient values ( $\beta$ ), 95% confidence limits (CL), and *p*-values are shown, marked with asterisks to indicate significance after controlling for multiple testing.

While the above results partially confirm our hypothesis that individuals should walk more briskly when the experimenter exerted additional effort and time to hold the box, the manner in which the box was held did not have a statistically significant effect. Specifically, when the box was picked up and held at waist level, participants approached the hand off at 1.69 x10^−2^ m s^−1^ per kg of box mass more quickly than if the box lay on the ground. In fact, when the box was picked up and held even higher, at chest level, with arms outstretched (presumably substantially more effort exerted for a large box), participants only approached the hand off at 1.37 x10^−2^ m s^−1^ per kg of box mass more quickly. However, the difference between these two effects was not significantly different (*p* = 0.31) and the AIC was slightly more negative (−1485.8 versus −1461.7) when the pose data were reduced to only two conditions for analysis: box held versus box on the ground. Because there was no evidence that participants meaningfully distinguished these conditions, subsequent analyses treated them as a single held condition to reduce model complexity.

### Participant experiment responses varied with individual characteristics

Participant responses during the experiment, quantified by coefficient *β₂* representing the box mass-pose interaction, varied substantially across individuals. When compared with composite empathy scores, *β₂* exhibited a very weak positive correlation, indicating that participants with higher empathy scores tended to demonstrate slightly larger experimental responses; however, this relationship was not statistically significant (*R* = 0.159, *p* = 0.640, N=11). Deeper inspection of the data revealed that this relationship may have been influenced by a small number of outlying observations, with the correlation increasing to *R* = 0.457 following removal of the largest outlier and to *R* = 0.664 following further removal of the next largest outlier (see Supplemental Information). However, due to low sample size and a lack of valid justification for removing outliers, we strongly discourage overinterpreting any potential relationship based on the current data alone.

Additional exploratory analyses suggested that participant responses may also vary with individual characteristics. On average, female participants exhibited lower experimental responses than male participants (*β*_2_ = 0.012 ± 0.005 vs. 0.019 ± 0.007 m s^−1^ kg^−1^, respectively), despite having similar empathy scores (49.3±7.0 vs. 48.6±8.8). Participant responses also tended to increase slightly with age (*R* = 0.388) and decrease with average walking speed (*R* = −0.479). However, none of the observed relationships reached statistical significance (*p* ≥ 0.083). Small sample size strongly limits any interpretation of this exploratory analysis but may instead point to future avenues of investigation. Individual data and characteristics may be viewed in the Supplemental Information associated with this manuscript for further inspection.

## Discussion

### Sensitivity to the energy and time costs of others

Optimal trade offs between energy and time costs have been proposed as an explanation of self-selected walking speed varying with distance. Consistent with this framework, participants in the present study exhibited the same qualitative behavior observed in previous studies, increasing average walking speed with walking distance and converging on an asymptotic speed at longer distances (Carlisle & Kuo, 2023; Seethapathi & Srinivasan, 2015). In the nominal condition, where participants approached a zero-load box resting on the ground, parameters of the speed-distance relationship were comparable. While the nominal speed asymptote of 1.25 m s^−1^ (CL: 1.18, 1.32 m s^−1^) was substantially lower than Carlisle & Kuo’s reported value of 1.52 m s^−1^, this is unsurprising given we report average approach speeds (including acceleration/deceleration phases) and they report peak speeds. Seethapathi & Srinivasan also report higher average speeds of about 1.38 m s^−1^, but notably, this was at a much farther walking distance of about 89 m. More broadly, our nominal speed asymptote of 1.25 m s^−1^ is largely consistent with speeds corresponding to the minimum cost of transport, as are commonly reported in laboratory settings (Browning et al., 2006; Martin et al., 1992; Ralston, 1958; Willis et al., 2005). With respect to the rate constant of the speed-distance curve, Carlisle & Kuo reported a value of 0.53 m^−1^ compared to 0.44 m^−1^ (0.42, 0.45 m^−1^) in the current study. Some sampling variance is to be expected given small sample sizes (N=11 or 10 across the comparison studies discussed here).

Beyond the nominal condition, where movement strategies were consistent with those that minimize energy and time costs, participants also seemed to respond to costs incurred by the experimenter acting on their behalf. When the experimenter lifted and held the box, participants often approached at a faster speed, in particular at farther walking distances and when the held box was more massive (e.g., 1.29 m s^−1^ vs. 1.19 m s^−1^ with a 6.8 kg box, *p* = 3.85 x 10^−6*^). We interpret these behavioral changes to reflect participant sensitivity to the energy and time costs of the experimenter spent helping the participant. In a literal sense, lifting and holding the box imposes both energy and time costs on the experimenter. Holding a heavier box for a longer duration requires greater sustained force and is expected to incur a greater energy cost related to the force-time integral of isometric muscle contractions (Crow & Kushmerick, 1982). Longer durations, associated with farther walking distances and slower approach speeds, also require the experimenter to spend more time waiting for the participant to reach them. Although lifting the box from the ground further requires mechanical work to increase the potential energy of the box, this cost cannot be reduced by arriving sooner, assuming the experimenter lifts the box regardless. Still, the act of lifting the box may help signal to the participant that some effort is being spent on their behalf. Both the time duration of holding the box and the associated force-time integral demands can be reduced when participants walk more quickly. Consistent with this interpretation, the nonlinear mixed-effects model revealed a significant interaction between box mass and hand off pose (*β*_2_ = 0.015 m s^−1^ kg^−1^, *p* = 3.85 x 10^−6^), indicating that the influence of holding the box became progressively larger as box mass increased.

Surprisingly, holding the box at the chest level was not associated with faster speed asymptotes than when the box was held at waist level (*p* = 0.31), despite additional mechanical work to lift the box higher and greater shoulder moments required to hold the box away from the body with arms outstretched. While we cannot fully account for this finding, one possible explanation is that participants were unable to accurately perceive the additional effort to hold the box away from the body. Since participants primarily viewed the holder in the frontal plane, visual cues indicating the anterior displacement of the box and the resulting increase in shoulder loading were likely limited.

It is possible that social facilitation may have resulted in increased walking speeds in the presence of the experimenter observing participants’ gait (Van Meurs et al., 2024). However, observation was maintained throughout the experiment, and so any such effects might apply to all walking data. As such, social facilitation likely does not explain statistical differences associated with box mass or hand off pose. Instead, the findings are broadly consistent with previous observations that individuals alter their behavior when another person incurs effort on their behalf, such as when a stranger holds a door open (Santamaria & Rosenbaum, 2011). However, few other studies have measured prosocial behaviors related to gait effort with relatively controlled experimental manipulations. The present study provides a quantitative framework for understanding these behavioral changes as a trade-off between one’s own energy and time costs and the perceived energy and time costs extracted from others, as mediated through an experimentally prescribed social interaction.

Importantly, any sensitivity to the energy costs of others must necessarily be inferred. Unlike an individual’s internal metabolic expenditure (Medrano et al., 2022; Selinger et al., 2015), the energy states of other individuals cannot be directly accessed, and even internal estimates of metabolic expenditure are sometimes insensitive (Wong et al., 2017), despite strong tendencies toward energy-optimal behaviors in other contexts (Selinger et al., 2015). Alternative theories sometimes consider the possibility that the central nervous system may combine proprioception and other signals to form a proxy of energy cost. For example, it is well know that internal models of force-based exertion are constructed from previous personal experiences, such as sensorimotor predictions of appropriate force control applied to external loads (Flanagan & Beltzner, 2000; Flanagan & Wing, 1997).

It is possible that internal models are extrapolated and projected onto the perceived experiences of others. Specifically, participants visually observing the experimenter lifting and holding a box may infer associated force-based costs from the size of the box as well as the time duration required for hand off. Furthermore, participants had previously handled the boxes personally and thus possessed direct experience regarding their mass and resulting weight. Together, these cues may have allowed participants to estimate the effort incurred by the experimenter and adjust their walking behavior accordingly. In fact, previous studies have shown that trained observers and coaches can accurately perceive exertion in children walking on an inclined treadmill (Robertson et al., 2006) and athletes during training sessions (Murphy et al., 2014; Wallace et al., 2009), respectively. There is also evidence showing that external ratings of perceived exertion are correlated with effort in tasks such as lifting boxes (Wangenheim et al., 1987) and other forceful exertion tasks such as pushing, pulling, lifting, and gripping (Bao et al., 2010).

### Individual characteristics and responses to the experiment

Although none of the relationships between participant characteristics and experimental response reached statistical significance, some exploratory trends were observed. Participants with slower nominal walking speeds tended to exhibit larger responses to the experimental manipulation, increasing their walking speeds to a greater extent when the experimenter held a large box (*R* = −0.479, *p* = 0.136). This observation is consistent with the proposed optimization framework. Since the energy cost of walking increases nonlinearly with speed (Bastien et al., 2005), the marginal cost of increasing walking speed is expected to be smaller for individuals who typically walk more slowly than for those who already walk at relatively high speeds. Consequently, naturally slower walkers may be more willing to increase their speed at a lower marginal cost, when doing so reduces the perceived energy and time experienced by another person.

Other modest and moderate relationships between the experiment response, empathy score, age, and sex were observed. Participants with higher empathy scores tended to exhibit slightly larger responses in the experiment, although this relationship was not statistically significant (*R* = 0.159, *p* = 0.640). The correlation coefficient did increase markedly following removal of the largest outlier (*R* = 0.457), and by even more after removing the second largest outlier (*R* = 0.664), but the correlation was still not found to be statistically significant (*p* = 5.12 x 10^−2^). Similarly, older participants tended to report higher empathy scores and exhibited somewhat larger experimental responses than younger participants (*R* = 0.388, *p* = 0.239). This trend is consistent with studies that show older adults are sometimes more willing than younger adults to exert effort in favor of prosocial behaviors after accounting for physical ability (Byrne et al., 2023; Lockwood et al., 2021). Finally, despite having similar empathy scores (males: 48.6±8.8; females: 49.3±7.0) male participants exhibited larger experimental responses than females, when approaching larger boxes over farther walking distances (*R* = 0.545, *p* = 0.083). Overall, the sample empathy scores of this study were comparable with previous values reported in healthy adult samples: mean ± s.d. = 49.00 ± 7.47 versus 44.54 ± 7.70, respectively (Spreng et al., 2009).

Given a limited sample size and the absence of statistical significance, these observations should be interpreted with caution. Numerous social, cultural, and individual factors may influence how people perceive and respond to assistance provided by others, and the present study was not designed to establish individual characteristics that determine experiment responses. At the same time, they perhaps point to potential trends obscured by Type II error and may be more credibly discerned in future studies with larger and more diverse samples, to help clarify whether demographic or psychosocial characteristics influence responsiveness to the energy and time costs experienced by others.

### Limitations

Several important limitations should be considered when interpreting the findings of this study. First, it is difficult to determine whether the observed effects were driven primarily by the social interaction itself or by differences in the biomechanical task performed at the endpoint. When the box was placed on the ground, participants were required to bend down and lift it, whereas when the box was held, the hand off occurred at approximately waist or chest height. To reduce the influence of these differences, only the approach phase was analyzed, with walking duration and distance measured from the first to the second to last step before reaching the box. Nevertheless, participants may have modified their approach behavior in anticipation of the subsequent biomechanical task. We believe this explanation is unlikely to fully account for the observed findings because the effect of hand off pose reversed across walking distances; participants approached more slowly when the box was held at short distances but more quickly when the box was held at long distances. Anticipation and transitioning to the endpoint task may account for a more hesitant approach speed at short walking distances, as the box was often still being raised to the appropriate height (waist or chest) even after the participant arrived for the hand off. As such, participants may have approached more slowly due to the uncertainty of the hand off pose (Krüger & Hermsdörfer, 2019), additional attentional resources allocated toward monitoring the hand off condition (Yogev-Seligmann et al., 2008), or motor interference involved with simultaneously coordinating the hand off with walking (Beurskens et al., 2016). Conversely, the largest increases in walking speed occurred during farther walking bouts, suggesting that the social interaction may have been the dominant influence under those conditions.

Although the findings of this study are consistent with the energy-time optimization framework, it is possible that participants responded to other biomechanical costs such as effort or other force-based costs (Morel et al., 2017; Pedotti et al., 1978; Rebula & Kuo, 2015). It can be difficult to distinguish between the effects of mechanical energy and force due to their strong association, and this study cannot provide such distinctions. On the other hand, walking speed has previously been explained through an optimal trade off between energy and time, and so the current findings are interpreted likewise.

Another limitation is the small sample size of the experiment (N=11). Although several relationships were observed between experimental response and participant characteristics, none reached statistical significance. As a result, it remains unclear whether the observed trends reflect genuine underlying relationships or simply sampling variability. It is therefore possible that some of the non-significant findings represent Type II errors that would become statistically significant in a larger sample. More broadly, the extent to which the present findings generalize to the wider population remains uncertain.

Generalizability is further limited by the highly controlled and contrived nature of the experimental task. Although the experiment was intended to be more ecological than many laboratory-based walking studies, participants were explicitly instructed to retrieve a box from the experimenter. Consequently, participants may have perceived themselves as already assisting the experimenter by completing the task, potentially reducing any motivation to further minimize the experimenter’s efforts. Furthermore, social responses likely differ substantially in real-world settings involving close friends, family members, coworkers, or strangers encountered in everyday situations. Future studies should examine whether the magnitude of the observed effects varies across different social contexts and interpersonal relationships.

Finally, the behavior examined in the present study is likely influenced by numerous social, psychological, and environmental factors. For example, participant responses to the empathy questionnaire may have been influenced by social desirability bias (Crowne & Marlowe, 1960). while behavior during the walking trials may have been shaped by social expectations or expectations of the experimental task itself. Data collection took place in the Human Performance Laboratory at the University of Calgary, a shared research space that occasionally experienced pedestrian traffic. Although no obvious effects of passersby were observed either qualitatively or quantitatively, it remains possible that social context influenced participant behavior. Similarly, responses may depend on characteristics of the individual providing assistance. In the present study, all trials involved the same experimenter, and participants may have responded differently to an individual of a different age, sex, gender, physical stature, ethnicity, or perceived social role. Consistent with this possibility, exploratory analyses suggested potential differences across participant age and sex, although none of these relationships were statistically significant. Future studies should explicitly investigate how characteristics of both the participant and the assisting individual influence responsiveness to the energy and time costs experienced by others.

## Conclusions

The present study examined the influence of a simple social interaction on self-selected walking speed. Consistent with previous work, participants adjusted their walking speeds according to walking distance in a manner consistent with optimal energy-time trade offs. However, walking behavior was also influenced by whether a box was held by another individual or left on the ground. Participants approached held boxes more slowly at short walking distances but more quickly at farther walking distances, with the effect becoming increasingly pronounced as box mass increased. Several participant characteristics exhibited modest-to-moderate associations with individual responses in the experiment, including empathy score, age, sex, and nominal walking speed. However, none of these relationships reached statistical significance, perhaps in part due to a limited sample size. Social interactions are rarely incorporated into laboratory studies of human walking despite their ubiquity in everyday life. The present findings demonstrate that social context can measurably influence self-selected walking behavior and suggest that incorporating social interactions into locomotion research may provide additional insight into how humans move and make decisions in real-world environments.

## Acknowledgements

We are grateful to the Human Performance Laboratory, the Department of Kinesiology, and to Jeremy Wong at the University of Calgary for facilitating the gait experiments described herein.

## Data and resource availability

All data and data processing scripts associated with this manuscript are available on Zenodo.org (Schroeder, 2026) at https://doi.org/10.5281/ZENODO.21459425. All other relevant data can be found within the article and its supplementary information.

## Competing Interests

No competing interests declared.

## Funding

This research received no specific grant from any funding agency in the public, commercial or not-for-profit sectors.

## List of Symbols

|  |  |  |
| --- | --- | --- |
| $A$ | asymptotic approach speed | (m s<sup>-1</sup>) |
| $A_0$ | nominal speed asymptote | (m s<sup>-1</sup>) |
| AIC | Akaike's Information Criterion |  |
| $B_1$ | individual effect of mass-pose interaction on speed in regression model | (m s<sup>-1</sup> kg<sup>-1</sup>) |
| $B_2$ | individual effect of individual speed asymptote in regression model | (m s<sup>-1</sup>) |
| CL | 95% confidence limits |  |
| $D$ | walking distance of approach | (m) |
| $k$ | rate constant describing speed-distance relationship | (m<sup>-1</sup>) |
| $k_0$ | nominal rate constant | (m<sup>-1</sup>) |
| $M$ | body mass | (kg) |
| $P$ | hand off pose | |
| RMSE | root mean square error | (m s<sup>-1</sup>) |
| $S$ | approach speed | (m s<sup>-1</sup>) |
| sd | standard deviation |  |
| $T$ | time duration of walking approach | (s) |
| $t$ | time | (s) |
| $v$ | step velocity | (m s<sup>-1</sup>) |
| $\beta_1$ | effect of mass on speed in nonlinear regression model | (m s<sup>-1</sup> kg<sup>-1</sup>) |
| $\beta_2$ | effect of mass-pose interaction on speed in regression model | (m s<sup>-1</sup> kg<sup>-1</sup>) |
| $\beta_3$ | effect of mass on rate constant in nonlinear regression model | (m<sup>-1</sup> kg<sup>-1</sup>) |
| $\beta_4$ | effect of mass-pose interaction on rate constant in regression model | (m<sup>-1</sup> kg<sup>-1</sup>) |

## Notes

https://doi.org/10.5281/ZENODO.21459425

### Competing Interest Statement

The authors have declared no competing interest.

## References

Akaike, H. (1974). A new look at the statistical model identification. IEEE Transactions on Automatic Control, 19(6), 716–723. 10.1109/TAC.1974.1100705

Bao, S., Howard, N., Spielholz, P., & Silverstein, B. (2010). Inter-observer reliability of forceful exertion analysis based on video-recordings. Ergonomics, 53(9), 1129–1139. 10.1080/00140139.2010.507879

Baron, R. S. (1986). Distraction-Conflict Theory: Progress and Problems. In Advances in Experimental Social Psychology (Vol. 19, pp. 1–40). Elsevier. 10.1016/S0065-2601(08)60211-7

Bastien, G. J., Willems, P. A., Schepens, B., & Heglund, N. C. (2005). Effect of load and speed on the energetic cost of human walking. European Journal of Applied Physiology, 94(1), 76–83. 10.1007/s00421-004-1286-z

Bertram, J. E. A. (2005). Constrained optimization in human walking: Cost minimization and gait plasticity. Journal of Experimental Biology, 208(6), 979–991. 10.1242/jeb.01498

Bertram, J. E. A., & Ruina, A. (2001). Multiple Walking Speed–frequency Relations are Predicted by Constrained Optimization. Journal of Theoretical Biology, 209(4), 445–453. 10.1006/jtbi.2001.2279

Beurskens, R., Steinberg, F., Antoniewicz, F., Wolff, W., & Granacher, U. (2016). Neural Correlates of Dual-Task Walking: Effects of Cognitive versus Motor Interference in Young Adults. Neural Plasticity, 2019, 1–9. 10.1155/2016/8032180

Browning, R. C., Baker, E. A., Herron, J. A., & Kram, R. (2006). Effects of obesity and sex on the energetic cost and preferred speed of walking. Journal of Applied Physiology, 100(2), 390–398. 10.1152/japplphysiol.00767.2005

Butki, B. D. (1994). Adaptation to effects of an audience during acquisition of rotary pursuit skill. Perceptual and Motor Skills, 79(3 Pt 1), 1151–1159.

Byrne, K. A., Lockwood, P. L., Ghaiumy Anaraky, R., & Liu, Y. (2023). Age Differences in Prosocial Behavior Depend on Effort Costs. The Journals of Gerontology: Series B, 78(6), 948–958. 10.1093/geronb/gbac194

Carlisle, R. E., & Kuo, A. D. (2023). Optimization of energy and time predicts dynamic speeds for human walking. eLife, 12, e81939. 10.7554/eLife.81939

Cornejo, C., Cuadros, Z., Morales, R., & Paredes, J. (2017). Interpersonal Coordination: Methods, Achievements, and Challenges. Frontiers in Psychology, 8, 1685. 10.3389/fpsyg.2017.01685

Crow, M. T., & Kushmerick, M. J. (1982). Chemical energetics of slow- and fast-twitch muscles of the mouse. Journal of General Physiology, 79(1), 147–166. 10.1085/jgp.79.1.147

Crowne, D. P., & Marlowe, D. (1960). A new scale of social desirability independent of psychopathology. Journal of Consulting Psychology, 24(4), 349–354. 10.1037/h0047358

Dunn, O. J. (1961). Multiple Comparisons among Means. Journal of the American Statistical Association, 56(293), 52–64. 10.1080/01621459.1961.10482090

Flanagan, J. R., & Beltzner, M. A. (2000). Independence of perceptual and sensorimotor predictions in the size–weight illusion. Nature Neuroscience, 3(7), 737–741. 10.1038/76701

Flanagan, J. R., & Wing, A. M. (1997). The Role of Internal Models in Motion Planning and Control: Evidence from Grip Force Adjustments during Movements of Hand-Held Loads. The Journal of Neuroscience, 17(4), 1519–1528. 10.1523/JNEUROSCI.17-04-01519.1997

Gutmann, A. K., Jacobi, B., Butcher, M. T., & Bertram, J. E. A. (2006). Constrained optimization in human running. Journal of Experimental Biology, 209(4), 622–632. 10.1242/jeb.02010

Hoyt, D. F., & Taylor, C. R. (1981). Gait and the energetics of locomotion in horses. Nature, 292(5820), 239–240. 10.1038/292239a0

Huber, M., Su, Y.-H., Krüger, M., Faschian, K., Glasauer, S., & Hermsdörfer, J. (2014). Adjustments of Speed and Path when Avoiding Collisions with Another Pedestrian. PLoS ONE, 9(2), e89589. 10.1371/journal.pone.0089589

Krendl, A., Gainsburg, I., & Ambady, N. (2012). The Effects of Stereotypes and Observer Pressure on Athletic Performance. Journal of Sport and Exercise Psychology, 34(1), 3–15. 10.1123/jsep.34.1.3

Krüger, M., & Hermsdörfer, J. (2019). Target Uncertainty During Motor Decision-Making: The Time Course of Movement Variability Reveals the Effect of Different Sources of Uncertainty on the Control of Reaching Movements. Frontiers in Psychology, 10, 41. 10.3389/fpsyg.2019.00041

Lockwood, P. L., Abdurahman, A., Gabay, A. S., Drew, D., Tamm, M., Husain, M., & Apps, M. A. J. (2021). Aging Increases Prosocial Motivation for Effort. Psychological Science, 32(5), 668–681. 10.1177/0956797620975781

Martin, P. E., Rothstein, D. E., & Larish, D. D. (1992). Effects of age and physical activity status on the speed-aerobic demand relationship of walking. Journal of Applied Physiology, 73(1), 200–206. 10.1152/jappl.1992.73.1.200

Medrano, R. L., Thomas, G. C., & Rouse, E. J. (2022). Can humans perceive the metabolic benefit provided by augmentative exoskeletons? Journal of NeuroEngineering and Rehabilitation, 19(1), 26. 10.1186/s12984-022-01002-w

Moussaïd, M., Perozo, N., Garnier, S., Helbing, D., & Theraulaz, G. (2010). The Walking Behaviour of Pedestrian Social Groups and Its Impact on Crowd Dynamics. PLoS ONE, 5(4), e10047. 10.1371/journal.pone.0010047

Murphy, A. P., Duffield, R., Kellett, A., & Reid, M. (2014). Comparison of Athlete–Coach Perceptions of Internal and External Load Markers for Elite Junior Tennis Training. International Journal of Sports Physiology and Performance, 9(5), 751–756. 10.1123/ijspp.2013-0364

Nessler, J. A., & Gilliland, S. J. (2009). Interpersonal synchronization during side by side treadmill walking is influenced by leg length differential and altered sensory feedback. Human Movement Science, 28(6), 772–785. 10.1016/j.humov.2009.04.007

Nicolas, A., & Hassan, F. H. (2023). Social groups in pedestrian crowds: Review of their influence on the dynamics and their modelling. Transportmetrica A: Transport Science, 19(1), 1970651. 10.1080/23249935.2021.1970651

Orendurff, M. S. (2008). How humans walk: Bout duration, steps per bout, and rest duration. The Journal of Rehabilitation Research and Development, 45(7), 1077–1090. 10.1682/JRRD.2007.11.0197

Paulus, P. B., & Cornelius, W. L. (1974). An Analysis of Gymnastic Performance under Conditions of Practice and Spectator Observation. Research Quarterly. American Alliance for Health, Physical Education and Recreation, 45(1), 56–63. 10.1080/10671188.1974.10615240

Ralston, H. J. (1958). Energy-speed relation and optimal speed during level walking. Internationale Zeitschrift Für Angewandte Physiologie Einschliesslich Arbeitsphysiologie, 17(4), 277–283. 10.1007/BF00698754

Rebula, J. R., Ojeda, L. V., Adamczyk, P. G., & Kuo, A. D. (2013). Measurement of foot placement and its variability with inertial sensors. Gait & Posture, 38(4), 974–980. 10.1016/j.gaitpost.2013.05.012

Robertson, R. J., Goss, F. L., Aaron, D. J., Tessmer, K. A., Gairola, A., Ghigiarelli, J. J., Kowallis, R. A., Thekkada, S., Liu, Y., Randall, C. R., & Weary, K. A. (2006). Observation of Perceived Exertion in Children Using the OMNI Pictorial Scale. Medicine & Science in Sports & Exercise, 38(1), 158–166. 10.1249/01.mss.0000190595.03402.66

Santamaria, J. P., & Rosenbaum, D. A. (2011). Etiquette and Effort: Holding Doors for Others. Psychological Science, 22(5), 584–588. 10.1177/0956797611406444

Schmidt-Nielsen, K. (1972). Locomotion: Energy Cost of Swimming, Flying, and Running. Science, 177(4045), 222–228. 10.1126/science.177.4045.222

Schroeder, R. (2026). Data supporting “Humans Modulate Walking Speed in Response to the Perceived Energy-Time Costs of Others” (Version Version 1.0) [Dataset]. Zenodo. 10.5281/ZENODO.21459425

Sebanz, N., Bekkering, H., & Knoblich, G. (2006). Joint action: Bodies and minds moving together. Trends in Cognitive Sciences, 10(2), 70–76. 10.1016/j.tics.2005.12.009

Seethapathi, N., & Srinivasan, M. (2015). The metabolic cost of changing walking speeds is significant, implies lower optimal speeds for shorter distances, and increases daily energy estimates. Biology Letters, 11(9), 20150486. 10.1098/rsbl.2015.0486

Selinger, J. C., O’Connor, S. M., Wong, J. D., & Donelan, J. M. (2015). Humans Can Continuously Optimize Energetic Cost during Walking. Current Biology, 25(18), 2452–2456. 10.1016/j.cub.2015.08.016

Sheridan, A., Marchant, D. C., Williams, E. L., Jones, H. S., Hewitt, P. A., & Sparks, A. (2019). Presence of Spotters Improves Bench Press Performance: A Deception Study. Journal of Strength and Conditioning Research, 33(7), 1755–1761. 10.1519/JSC.0000000000002285

Spreng, R. N., McKinnon, M. C., Mar, R. A., & Levine, B. (2009). The Toronto Empathy Questionnaire: Scale Development and Initial Validation of a Factor-Analytic Solution to Multiple Empathy Measures. Journal of Personality Assessment, 91(1), 62–71. 10.1080/00223890802484381

Sylos-Labini, F., d’Avella, A., Lacquaniti, F., & Ivanenko, Y. (2018). Human-Human Interaction Forces and Interlimb Coordination During Side-by-Side Walking With Hand Contact. Frontiers in Physiology, 9, 179. 10.3389/fphys.2018.00179

Tukey, J. W. (1977). Exploratory data analysis. Addison-Wesley Pub. Co.

Van Meurs, E., Greve, J., & Strauss, B. (2024). Moving in the presence of others – a systematic review and meta-analysis on social facilitation. International Review of Sport and Exercise Psychology, 17(2), 980–1012. 10.1080/1750984X.2022.2111663

Wallace, L. K., Slattery, K. M., & Coutts, A. J. (2009). The Ecological Validity and Application of the Session-RPE Method for Quantifying Training Loads in Swimming. Journal of Strength and Conditioning Research, 23(1), 33–38. 10.1519/JSC.0b013e3181874512

Wangenheim, M., Borg, G., & Holzmann, P. (1987). A Psychophysical Study of Work-Related Stress Using Observer Ratings. Upsala Journal of Medical Sciences, 92(1), 1–17. 10.3109/03009738709178674

Willis, W. T., Ganley, K. J., & Herman, R. M. (2005). Fuel oxidation during human walking. Metabolism, 54(6), 793–799. 10.1016/j.metabol.2005.01.024

Wong, J. D., O’Connor, S. M., Selinger, J. C., & Donelan, J. M. (2017). Contribution of blood oxygen and carbon dioxide sensing to the energetic optimization of human walking. Journal of Neurophysiology, 118(2), 1425–1433. 10.1152/jn.00195.2017

Worringham, C. J., & Messick, D. M. (1983). Social Facilitation of Running: An Unobtrusive Study. The Journal of Social Psychology, 121(1), 23–29. 10.1080/00224545.1983.9924462

Yogev-Seligmann, G., Hausdorff, J. M., & Giladi, N. (2008). The role of executive function and attention in gait. Movement Disorders, 23(3), 329–342. 10.1002/mds.21720

Zivotofsky, A. Z., & Hausdorff, J. M. (2007). The sensory feedback mechanisms enabling couples to walk synchronously: An initial investigation. Journal of NeuroEngineering and Rehabilitation, 4(1), 28. 10.1186/1743-0003-4-28

