## Supplementary Information for "Humans Modulate Walking Speed in Response to the Perceived Energy-Time Costs of Others"

**Table S1**

Summary of Participant Characteristics

| Sex | Age (yrs) | Body Mass (kg) | Height (m) | Empathy Score (/64) | Box-Pose Effect, $\beta_2 + B_1$ (m s$^{-1}$ kg$^{-1}$) | Nominal speed, $A_0 + B_2$ (m s$^{-1}$) |
|---|---|---|---|---|---|---|
| Female | 23 | 61.2 | 1.75 | 51 | 0.010 | 1.452 |
| Female | 21 | 83.0 | 1.70 | 52 | 0.010 | 1.210 |
| Female | 41 | 62.1 | 1.62 | 56 | 0.021 | 1.309 |
| Female | 27 | 66.7 | 1.62 | 48 | 0.008 | 1.337 |
| Female | 21 | 62.1 | 1.65 | 36 | 0.006 | 1.360 |
| Female | 38 | 79.8 | 1.79 | 53 | 0.015 | 1.171 |
| Male | 26 | 122.4 | 1.83 | 44 | 0.016 | 1.045 |
| Male | 20 | 68.9 | 1.88 | 39 | 0.017 | 1.296 |
| Male | 30 | 91.1 | 1.50 | 45 | 0.031 | 1.107 |
| Male | 26 | 79.8 | 1.87 | 54 | 0.012 | 1.169 |
| Male | 20 | 78.9 | 1.78 | 61 | 0.019 | 1.302 |

Individual participant characteristics are listed including Sex, Age, Body Mass, and Height. The cumulative Empathy Score is also displayed (out of 64), where higher scores indicate higher levels of self-reported emotional empathy. Additionally, individual responses to the experiment are indicated as the coefficient on the interaction effect of box mass with hand off pose, where $\beta_2$ is the sample coefficient (fixed effect) and $B_1$ is the individual coefficient (random effect) from the nonlinear regression model (see Table 1, Eqs. 1-3 in the main text). Finally, Nominal Speed is listed, indicating the speed asymptote when approaching a zero-load box sitting on the ground, where $A_0$ is the sample parameter (fixed effect) and $B_2$is the individual parameter (random effect) from the nonlinear regression model.

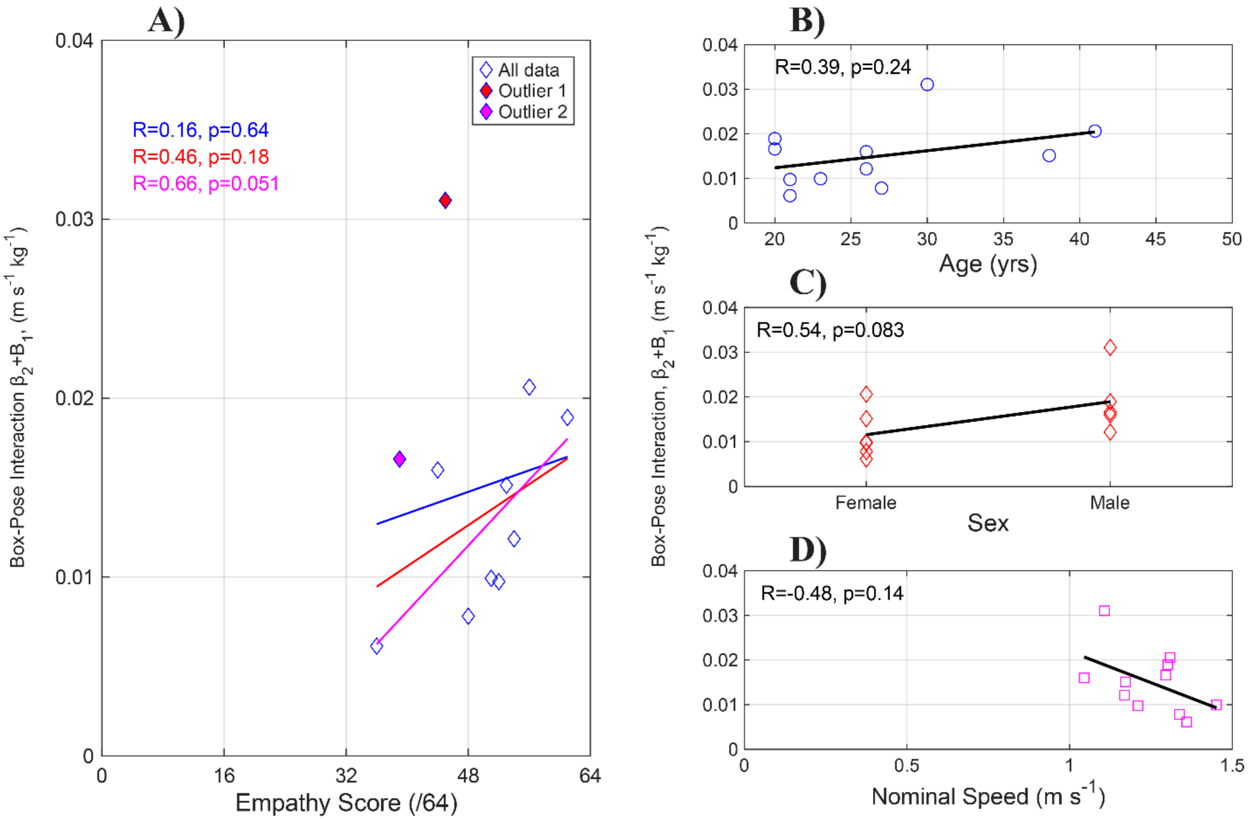


**Figure S1.** Relationships to individual participant characteristics. **(A)** Individual cumulative Empathy Score (out of 64) is evaluated for its effect on the Box-Pose Interaction ($\beta_2 + B_1$) in the regression model – a quantification of the primary experimental response of individuals. Linear regressions, correlation coefficients, and *p*-values are shown for all data (blue), all data minus the largest outlier (red), and all data minus the two largest outliers (magenta). **(B)** The effect of participant Age, **(C)** Sex, and **(D)** Nominal Speed on the Box-Pose Interaction are shown with linear regression lines, correlation coefficients and corresponding *p*-values.
